# Genetic footprint of population fragmentation and contemporary collapse in a freshwater cetacean

**DOI:** 10.1101/094995

**Authors:** Minmin Chen, Michael C. Fontaine, Yacine Ben Chehida, Jinsong Zheng, Frédéric Labbe, Zhigang Mei, Yujiang Hao, Kexiong Wang, Min Wu, Qingzhong Zhao, Ding Wang

## Abstract

Understanding demographic trends and patterns of gene flow in an endangered species is crucial for devising conservation strategies. Here, we examined the extent of population structure and recent evolution of the critically endangered Yangtze finless porpoise (*Neophocaena asiaeorientalis asiaeorientalis*). By analysing genetic variation at the mitochondrial and nuclear microsatellite loci for 148 individuals, we identified three populations along the Yangtze River, each one connected to a group of admixed ancestry. Each population displayed extremely low genetic diversity, consistent with extremely small effective size (≤92 individuals). Habitat degradation and distribution gaps correlated with highly asymmetric gene-flow that was inefficient in maintaining connectivity between populations. Genetic inferences of historical demography revealed that the populations in the Yangtze descended from a small number of founders colonizing the river from the sea during the last Ice Age. The colonization was followed by a rapid population split during the last millennium predating the Chinese Modern Economy Development. However, genetic diversity showed a clear footprint of population contraction over the last 50 years leaving only ~2% of the pre-collapsed size, consistent with the population collapses reported from field studies. This genetic perspective provides background information for devising mitigation strategies to prevent this species from extinction.

## Introduction

Dispersal and gene flow in a meta-population maintain local demographic and genetic variation, thus increasing the probability of species persistence^1,2^. Persistence of wide-ranging animals occupying fragmented landscapes depends on the matrix quality of the habitat and the ability of individuals to move among habitat patches^3^, and corridors facilitating this movement across such landscape^4–6^. Along the Yangtze River (China, Fig. 1), anthropogenic activities of the past 50 years have put intense pressure on the freshwater ecosystem leading to habitat degradation, species range fragmentation and extinction of some emblematic endemic species, such as the Yangtze River dolphin or baiji (*Lipotes vexillifer*)^7^. Today, the Yangtze finless porpoise (*Neophocaena asiaeorientalis asiaeorientalis* or YFP) has become the only surviving freshwater cetacean now found in China and the world’s only freshwater porpoise species^8^.

**Figure 1.**
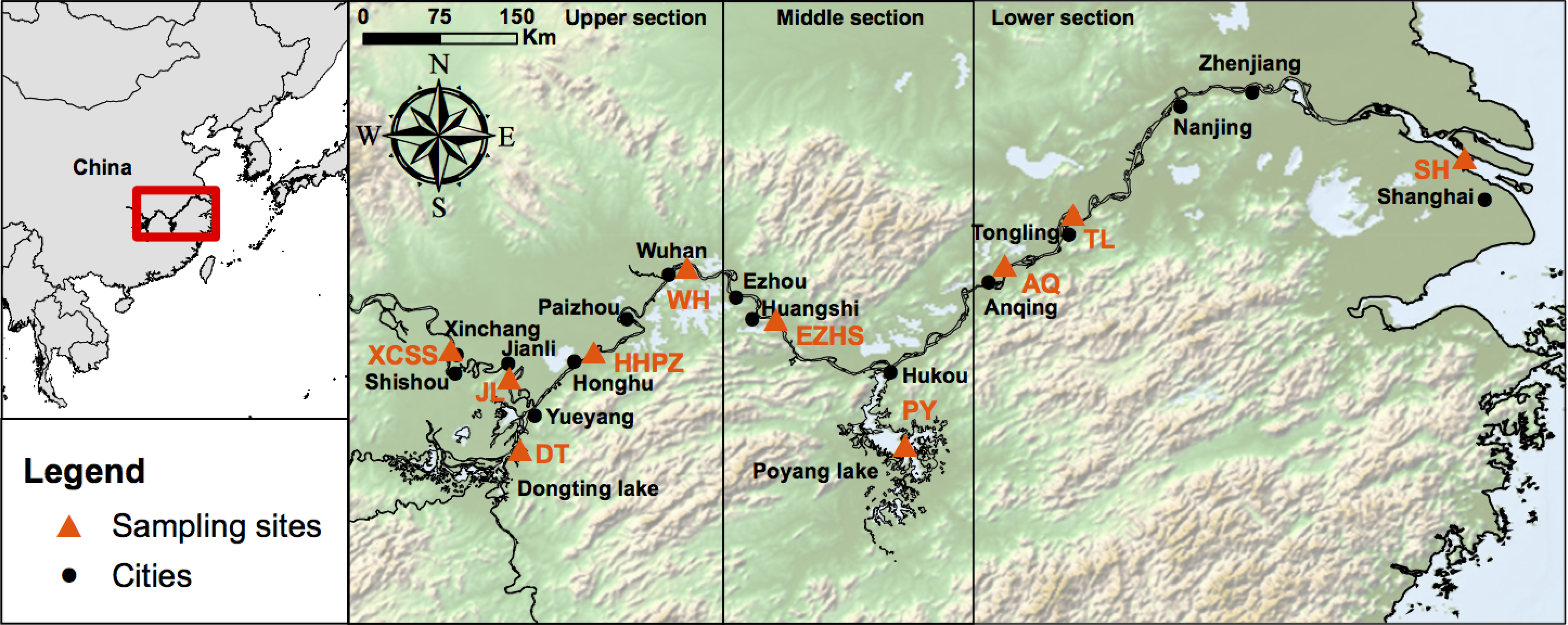
Maps showing the sampling distribution of the Yangtze finless porpoises in the Yangtze River. The top-left insert shows the location of the studied area highlighted by a red rectangle. On the right, the map shows the sampling locations (orange triangles) and their acronyms based on the neighbouring cities. Figure created using ArcGIS 10.3 software using the open source data from the ETOPO1 Global Relief Model^63^ (https://www.ngdc.noaa.gov/mgg/global/).

Endemic to the Yangtze River drainage (Fig. 1), the YFP is now primarily restricted to the middle-lower Yangtze Channel and two large appended lakes: Dongting Lake (DT) and Poyang Lake (PY)^9^. The subspecies has occasionally been reported from some of the larger adjacent tributaries though this is now rare^10–12^. The amount of river and lake habitat available to this subspecies is relatively small compared to that available to marine populations of finless porpoises, which occur in coastal waters from Japan to the Arabian Sea^8^. However, YFP abundance has suffered from dramatic reductions from 2,500 individuals in 1991^10^ to 1,800 in the end of 2006, as estimated by the Yangtze Freshwater Dolphin Expedition in 2006 (YFDE2006)^13^. More recent surveys conducted during the Yangtze Freshwater Dolphin Expedition conducted in 2012 (YFDE2012) reported that populations declined even further to ~1,040 individuals, including ~500 porpoises in the Yangtze main stream, ~450 in PY and ~90 in DT ^14^ With such rapid range contraction, Mei *et al*.^15^ estimated that the YFP may become extinct within the next 60 years or less^14^. The YFP was thus recently reclassified as a Critically Endangered sub-species on the IUCN Red List^9^. As a top predator, the survival of the finless porpoise depends heavily on habitat suitability, food availability and maintenance of corridors allowing dispersal between populations. However, with increasing underwater noise from boat traffic and incidental catches in fisheries, food and habitat resources for the species have become increasingly scattered and fragmented, and corridors across the landscape have been compromised by the booming of the Chinese economy over the last decades^13^. Despite these imminent threats, we still do not know how reduction in suitable habitats in the Yangtze River has reduced the number of breeding porpoises and how habitat fragmentation has impacted connectivity between populations and the population structure itself. This information is extremely difficult to quantify using direct observations. On the other hand, population genetic approaches can provide key insights about the current population structure and connectivity and historical population demography by leaving detectable footprints on the genetic diversity and its geographic structure^16,17^.

Previous phylogeographic analyses based on the non-coding mitochondrial Control Region (mtDNA-CR) of the finless porpoises from Chinese and Japanese waters documented evidence of a demographic expansion following the Last Glacial Maximum (LGM, ~24,000 yrs BP) and the colonization of Yangtze River from the Yellow Sea ~22,000 yrs BP ago^18,19^. A subsequent study within the Yangtze River by Chen et al.^20^ used mtDNA-CR and nuclear microsatellite loci and revealed evidence of genetic subdivisions within the YFP populations suggestive of population fragmentation. However, the fine scale population structure remained unclear and the processes shaping the genetic variation in the YFP unknown. Changes in connectivity between populations, dynamics of population expansion-contraction and demographic history are potent factors shaping the genetic variation in the YFP, but these were not investigated in further details so far.

In this study, we address these above questions by re-analysing the previously published data set of Chen et al.^20^ that combined fast evolving microsatellite loci together with more slowly evolving sequences from the mtDNA-CR. Specifically, in contrast with previous studies, we used a statistical population genetic framework in order to (i) resolve the fine-scale population structure using a combination of individual-based multivariate and Bayesian clustering approaches designed for weak genetic structure when it exists; (ii) estimate past and contemporaneous effective population sizes and connectivity among populations; and (iii) reconstruct the demographic history best fitting with the genetic diversity observed in the populations of the YFP using a quantitative model-based inferential population genetic framework relying on Approximate Bayesian Computation (ABC) approaches ^16,17,21–23^. Determining and quantifying to which extent populations in the YFP are fragmenting, loosing connectivity, and the magnitudes of the demographic trends are critical knowledge for designing conservation and management plans. For example, we still do not understand whether population fragmentation and decline have been triggered only recently by human activities during the past 50 years or if these patterns were initiated earlier by more long-term ecological and evolutionary processes and exacerbated lately during the Anthropocene.

## Results

**Genetic structure and diversity of the Yangtze finless porpoises.** The final data set consisted of 148 individuals sampled along the Yangtze River and the two main lakes (Dongting Lake (DT) and Poyang Lake (PY), Fig. 1) genotyped for 11 microsatellite loci and sequenced for a 597 base-pairs fragment of the hyper-variable region 1 (see Table 1 and materials and methods). The clustering of the microsatellite data using the Bayesian clustering algorithm of *STRUCTURE*^24–26^ provided consistent results over 10 replicated runs performed for each number *K* of cluster tested (Fig. 2a). The probability of the data greatly increased when two genetic clusters were modelled instead of one and showed the highest values on average over 10 replicates (Fig. S1). At K>2, the increase in probability decreased in average; however some runs displayed the highest probability of all runs at K=3 and 4 (Fig. S1). A visual inspection of the individual clustering for each K value (Fig. 2a) revealed that porpoises from the PY split from the other individuals of the Yangtze river at K=2, suggesting that these porpoises are highly differentiated from the others. When higher K values were tested (K=3 and 4) porpoises from PY, XCSS and TL localities were all identified as differentiated genetic units, while the porpoises from the in-between regions consisted of an admixed group sharing genetic ancestry with the three other populations (Fig. 2a and 2d). The three individuals at the mouth of the Yangtze River close to Shanghai city (SH) also seemed to depart from the other groups at K=4, but the low sample size (n=3) preclude any definitive conclusions. No further subdivision was observed beyond K=4 (result not shown).

**Figure 2.**
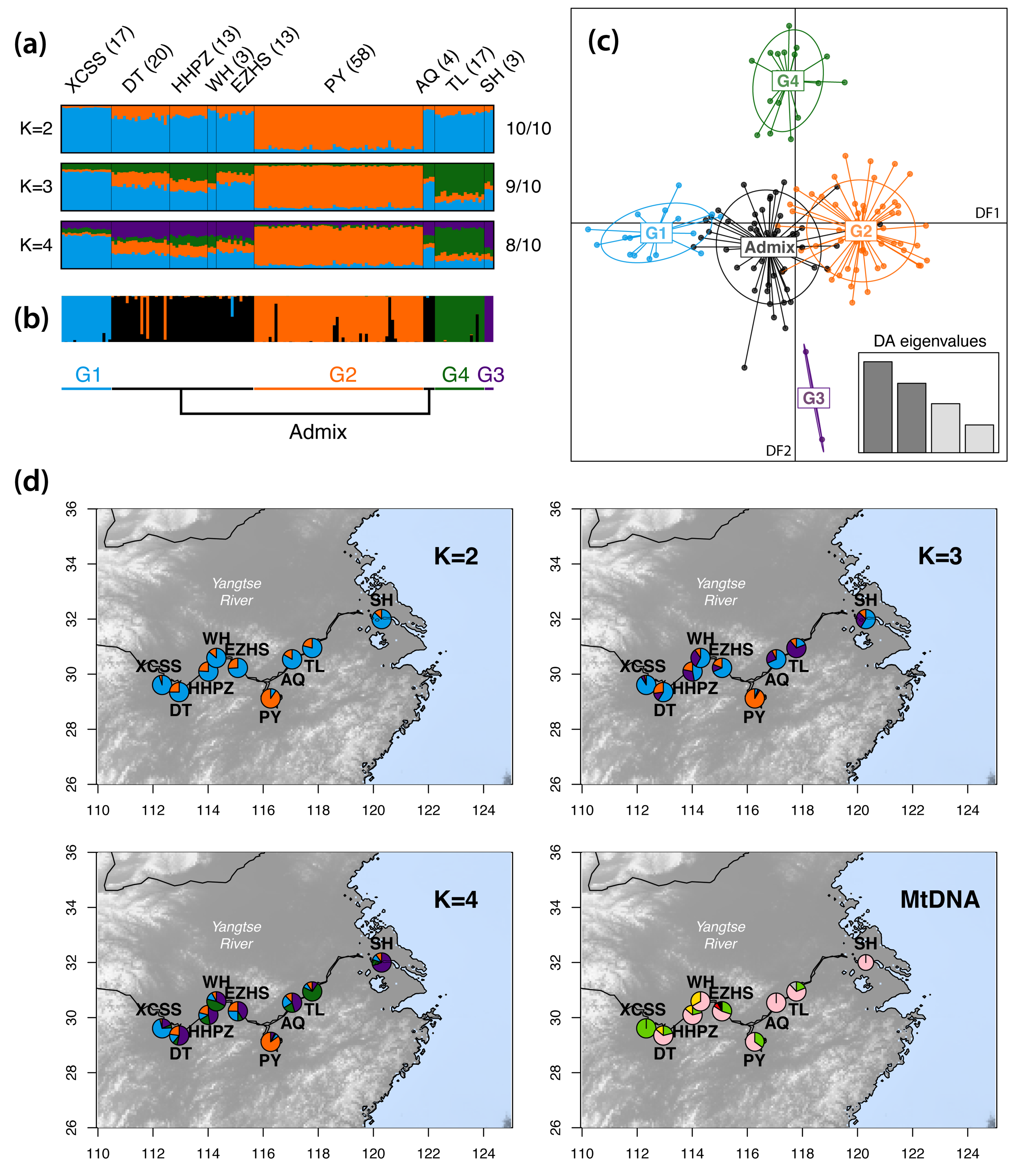
(a) Population structure estimated using the Bayesian clustering approach of *STRUCTURE*. Each individual is represented by a vertical line divided into K segments showing the admixture proportions from each cluster. Sample size in each locality is shown between brackets. Numbers on the right side of the barplot show the number of time this result was found out of the 10 replicates. (b) *DAPC* cluster membership probability plot of the 148 individuals. (c) Scatter plot showing the first two discriminant functions (DFs) of the *DAPC*. (d) Geographical distribution of the *STRUCTURE* admixture proportions and mitochondrial haplotype frequencies per localities. The mtDNA map is modified from Chen et al.^20^. Panel (a) was created using CLUMPAK^59^, panel (b) and (c) using R v.3.4.0^66^, and panel (d) using R v.3.4.0^66^, the package MARMAP v.0.9.5^62^, and the open source ETOPO1 Global Relief Model^63^ (https://www.ngdc.noaa.gov/mgg/global/).

**Table 1.** Genetic variation at the 11 microsatellites and mtDNA control region loci for each distinct population inferred from the *STRUCTURE* analysis.

|  | Total | PY | TL | XCSS | Admix |
| --- | --- | --- | --- | --- | --- |
| Microsatellite loci |  |  |  |  |  |
| $N_{Mic.} \pm SD$ (max) | 148 | 57.2 ± 0.32 (58) | 16.0 ± 0.4 (17) | 16.4 ± 0.36 (17) | 48.1 ± 1.3 (53) |
| NA | - | 1.4% | 4.8% | 3.7% | 9.3% |
| $A_r \pm SE$ | - | 4.82 ± 0.40 | 5.19 ± 0.52 | 5.25 ± 0.46 | 5.93 ± 0.45 |
| $pA \pm SE$ | - | 0.38 ± 0.09 | 0.55 ± 0.21 | 0.48 ± 0.12 | 0.69 ± 0.18 |
| $H_o \pm SD$ | - | 0.60 ± 0.06 | 0.70 ± 0.047 | 0.69 ± 0.05 | 0.72 ± 0.04 |
| $H_e \pm SD$ | - | 0.62 ± 0.05 | 0.62 ± 0.05 | 0.63 ± 0.04 | 0.68 ± 0.03 |
| $F_{IS} \pm SD$ | - | 0.041 ± 0.03<sup>ns</sup> | -0.123 ± 0.06<sup>ns</sup> | -0.112 ± 0.03<sup>ns</sup> | -0.057 ± 0.03<sup>ns</sup> |
| $M_{GW}^{\dagger}$ | - | 0.51*** | 0.46** | 0.44** | 0.53** |
| MtDNA control region |  |  |  |  |  |
| $N_{mtDNA}$ | 129 | 56 | 17 | 16 | 37 |
| $S$ | 7 | 2 | 1 | 0 | 5 |
| *Singleton* | 4 | 0 | 0 | 0 | 3 |
| *Shared P.* | 3 | 2 | 1 | 0 | 2 |
| *#hap.* | 7 | 3 | 2 | 1 | 5 |
| $Hd$ | 0.57 | 0.53 | 0.22 | 0 | 0.62 |
| $\pi$ (per site, %) | 0.112 | 0.096 | 0.037 | 0 | 0.150 |
| $\theta_W$ (per site, %) | 0.216 | 0.073 | 0.050 | 0 | 0.201 |
| $D^{\ddagger}$ | -1.08<sup>ns</sup> | 0.54<sup>ns</sup> | -0.49<sup>ns</sup> | 0 | -0.64<sup>ns</sup> |

$N_{Mic.}$ , microsatellite average sample size (max); NA: average proportion of missing data per locus, $A_r$ : allelic richness (estimated for a sample size of 26 individuals); $pA$ : Private allelic richness (estimated for a sample size of 3 individuals); $H_o$ and $H_e$ : observed and expected heterozygosity; $F_{IS}$ : Inbreeding coefficient; $M_{GW}$ , M of Garza and Williamson<sup>34</sup>; $N_{mtDNA}$ , MtDNA sample size; $S$ , number of segregating sites; *Singleton*: rare mutation observed only in one sequence among all; *Share P.*, shared polymorphism (mutation observed in at least two or more sequences) also known as parsimony informative site; *#hap.*, number of haplotypes; $Hd$ , haplotype diversity; $\pi$ , nucleotide diversity; $\theta_W$ , theta from $S$ or Theta-Watterson; $D$ , Tajima's $D$ . <sup>†</sup> The significance level of the $M_{GW}$ statistic was evaluated in DIYABC<sup>78</sup> using $1 \times 10^6$ coalescent simulations under a scenario of constant effective population size. The P-value indicate the proportion of simulations which provide a value below the observed one. <sup>‡</sup> The significance of $D$ values was estimated using 10 000 coalescent simulations in DNAsp<sup>73</sup>. ns: not significant (p-value > 0.05); \* p-value ≤ 0.05; \*\* p-value ≤ 0.01; \*\*\* p-value ≤ 0.001

The results from *STRUCTURE* were further validated using two multivariate approaches that do not rely on model assumptions^27^: a Discriminant Analysis of Principal Components (*DAPC*)^28^ and a Principal Component Analysis (PCA)^27,29^. Both methods showed a genetic structure consistent with the results of *STRUCTURE* (Fig. 2b, 2c and Fig. S2). The *DAPC* provided a clear-cut discrimination of the four groups identify by *STRUCTURE* (Fig. 2c). The first discriminant function (DF) discriminated XCSS and PY and the second DF TL and SH. The other porpoises recognized as admixed in *STRUCTURE* were located at the intersection of the four other groups. The proportions of successful reassignment (based on the discriminant functions) of individuals to their original clusters was high (Fig. 2b): 100% for XCSS and SH, 94.8% for PY, 94.1% for TL, and 94.3% for the admixed porpoises. These large values indicate clear-cut clusters. Finally, the PCA provided a similar picture as the *DAPC* and *STRUCTURE*, but with more overlap among the groups (Fig. S2).

None of the identified populations displayed significant departures from Hardy–Weinberg expectations as shown by the *F_IS_* values (Table 1). Porpoises from the admixed and TL groups showed the highest level of microsatellite genetic diversities and XCSS and PY the lowest, as estimated with the values of allelic richness (*Ar*), private allelic richness (*pA*) and expected heterozygosity (*H_e_*) (Table 1 and Fig. S3). Only the two extremes groups, PY and Admix, showed a significant difference in *A_r_* and *H_e_* (Wilcoxon signed-ranked test p-value < 0.05). The mitochondrial genetic diversity followed a similar trend with the admixed group showing the highest haplotype and nucleotide diversity followed by TL and PY (Table 1 and Fig. 2d). We found only one haplotype fixed in the XCSS group.

All populations showed significant differences in allelic frequencies for the microsatellite loci with *F_ST_* values ranging from 0.023 to 0.070 (Table 2). Notably *F_ST_* values between XCSS, PY and TL were relatively high (>0.05), while *F_ST_* values were intermediate between the Admix group and each distinct population (between 0.02 and 0.03, Table 2). For the mitochondrial locus (Table 2), *F_ST_* values were all significant except one (Admix vs. TL), and were especially strong between XCSS and all other groups, due to the fact that one haplotype is fixed in this population (Fig. 2d).

**Table 2.** Genetic differentiation between populations identified by Structure. Below the diagonal, pairwise *F_ST_* values and their 95% CI for microsatellite loci are provided as well as their associated P-value. Above the diagonal, *F_ST_* values are provided for the mtDNA locus with its corresponding P-value.

| $F_{ST-mic}$ \ $F_{ST-mtDNA}$ | XCSS | PY | TL | Admix |
| --- | --- | --- | --- | --- |
| XCSS | — | 0.590*** | 0.875*** | 0.446*** |
| PY | 0.070***<br>[0.038-0.104] | — | 0.109* | 0.011** |
| TL | 0.052***<br>[0.034-0.070] | 0.050***<br>[0.024-0.075] | — | 0.131<sup>ns</sup> |
| Admix | 0.029***<br>[0.011-0.047] | 0.023***<br>[0.014-0.032] | 0.023***<br>[0.010-0.037] | — |

ns: not significant (p-value > 0.05); \* p-value ≤ 0.05; \*\* p-value ≤ 0.01; \*\*\* p-value ≤ 0.001

**Contemporary effective population sizes and migration rates.** We used two methods for estimating contemporary effective population size (*Ne*) in each population from the microsatellite data: *NeEstimator*^30^ based on linkage disequilibrium among loci within population and *ONESAMP^31^* relying on an Approximate Bayesian Computation (ABC) approach. Both approaches provided comparable estimates of *Ne* for each population (Fig. 3 and Table 3). All estimated values were very low (<92 individuals), with the Admix and PY groups displaying the highest estimates, followed by TL and XCSS.

**Figure 3.**
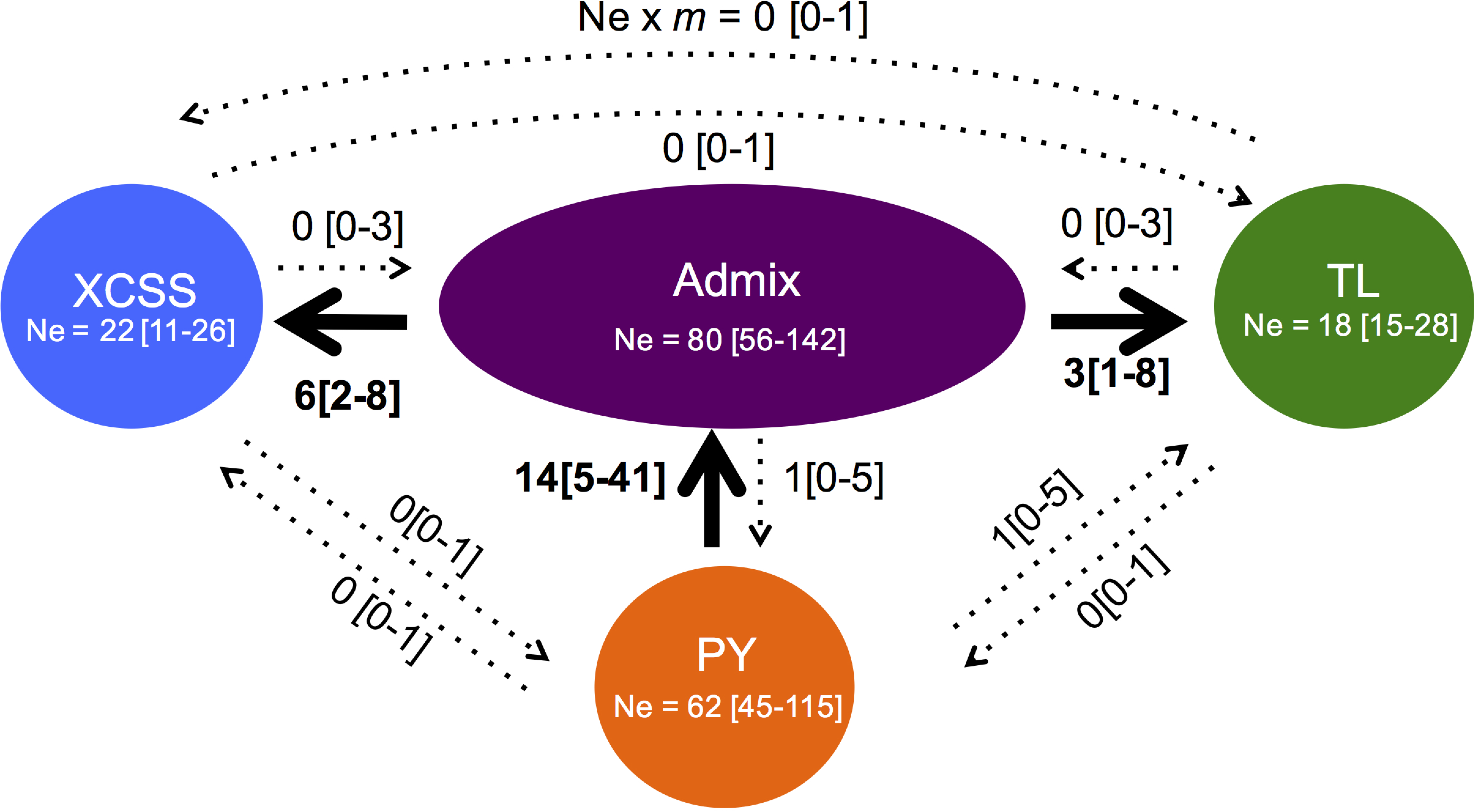
Recent gene flow (*Ne* × *m*) between populations estimated from *ONESAMP* and *BayesAss*. Confidence intervals are shown between squared brackets. Arrows show the effective migration rate significantly (plain) and not significantly different (dashed) from *Ne* estimates of *ONESAMP* (mean [95%CI]) in each population are provided in the circles.

**Table 3.**
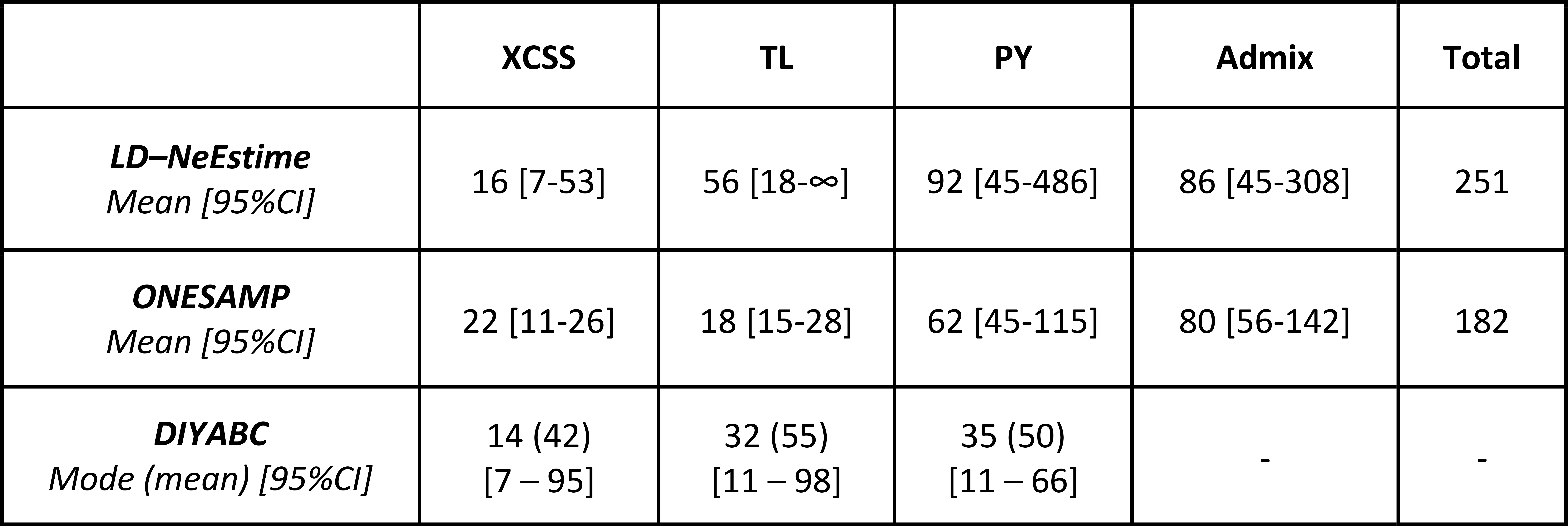
Effective population size (*Ne*) estimated in each population of the Yangtze finless porpoise. Values have been calculated using an estimator based on linkage disequilibrium between loci in *NeEstimator*^30^, using an ABC approach in *ONESAMP*^31^, and using DIYABC under SC2

We estimated migration rates (*m*) between pairs of groups over the last few generations using *BayesAss*^32^. Out of the 12 repeat runs, 10 showed good mixing properties with Bayesian Deviance values^33^ close to each other (mean ± SD: 8,915.28 ± 0.33) and convergent estimates for each parameter (see the material and methods). We thus combined them all to estimate the parameter values (Table S2). The number of effective migrants (*Ne* × *m*) per generation over the last generations were obtained by combining *Ne* estimates from *ONESAMP*^31^ with *m* estimates of *BayesAss*^32^ (Fig. 3). The three populations – TL, XCSS and PY – did not show any evidence of recent migration among each other, as indicated by the lower bound of the 95% highest probability density intervals (HPD) interval equal to 0 (Fig. 3 and Table S1). However, each one is connected to the Admix group with highly asymmetric gene flow. We detected significant unidirectional gene flow from PY to Admix and from Admix to TL and XCSS. Estimated *Ne × m* values from the Admix group to TL or XCSS are about the half of those from PY to the Admix group.

**Population demographic history.** Departure of the allelic or haplotypic frequency spectrum of microsatellite and mtDNA loci, respectively, from those expected under a scenario of constant population size can provide evidence of population size change. For microsatellite loci, we used the Garza and Williamson *M_GW_* ratio of the number of alleles to the range in allele size to detect evidence of population size contraction^34^. Genetic diversity at the microsatellite markers showed significant evidence of *Ne* contraction in each population, as suggested by the very small ratio values of the *M_GW_* statistic (Table 1). The *M_GW_* value estimated in each population was significantly smaller than expected under the assumption of constant population size (Table 1). In contrast, the Tajima’s D^35^ values (Table 1) estimated from the mtDNA-CR sequences in each population did not show any significant departure from the constant population size hypothesis.

We investigated further the demographic history best fitting with the genetic diversity of the combined microsatellite and mtDNA markers observed in the three distinct populations of the YFP using a coalescent-based ABC approach^22^. See the material and methods, appendix S1, Fig. S4, Table S2 and S3 for further details on the methodology. The first step in the ABC analysis was to identify the population branching order that best fit with the data. Out of the 10 scenarios tested (Fig. 4a), the ABC analysis showed that the trichotomy (SC1), which assumes that XCSS, PY and TL populations diverged at the same time, received the highest support. Indeed, the two distinct model choice approaches – a “standard” model choice procedure relying a logistic regression of Linear Discriminant Analysis (ABC-LDA) on the summary statistics^36^ and the recently developed Random Forest machine learning classification approach^37,38^ (ABC-RF) – identify this scenario SC1 as the best one with a posterior probability respectively of 66.7% with a 95%CI not overlapping with any other scenarios and 55.8 ± 2.5% (Fig. S5a and Table S4). This contrasted with the nine other models where the posterior probabilities estimated using the ABC-LDA were each lower than 8%. The simulation-based performance analysis of this ABC step (Table S4, S5 and Fig. S5) shows that 61.4% of the simulated dataset under this SC1 were correctly identified using the ABC-LDA procedure, leading to an average Type-I error rate (false negative) of 4.3% ranging from 1.7% to 9.7% and a total prior error rate *(i.e.* the average misclassification error)^37,38^ of 38.6% (62.9% using the ABC-RF). Simulations under the nine other competing scenarios led to a Type-II error rate of <12.9% incorrect assignment to SC1 (false positives) and a power (87.1%) to discriminate the best scenario from the others (Table S4).

**Figure 4.**
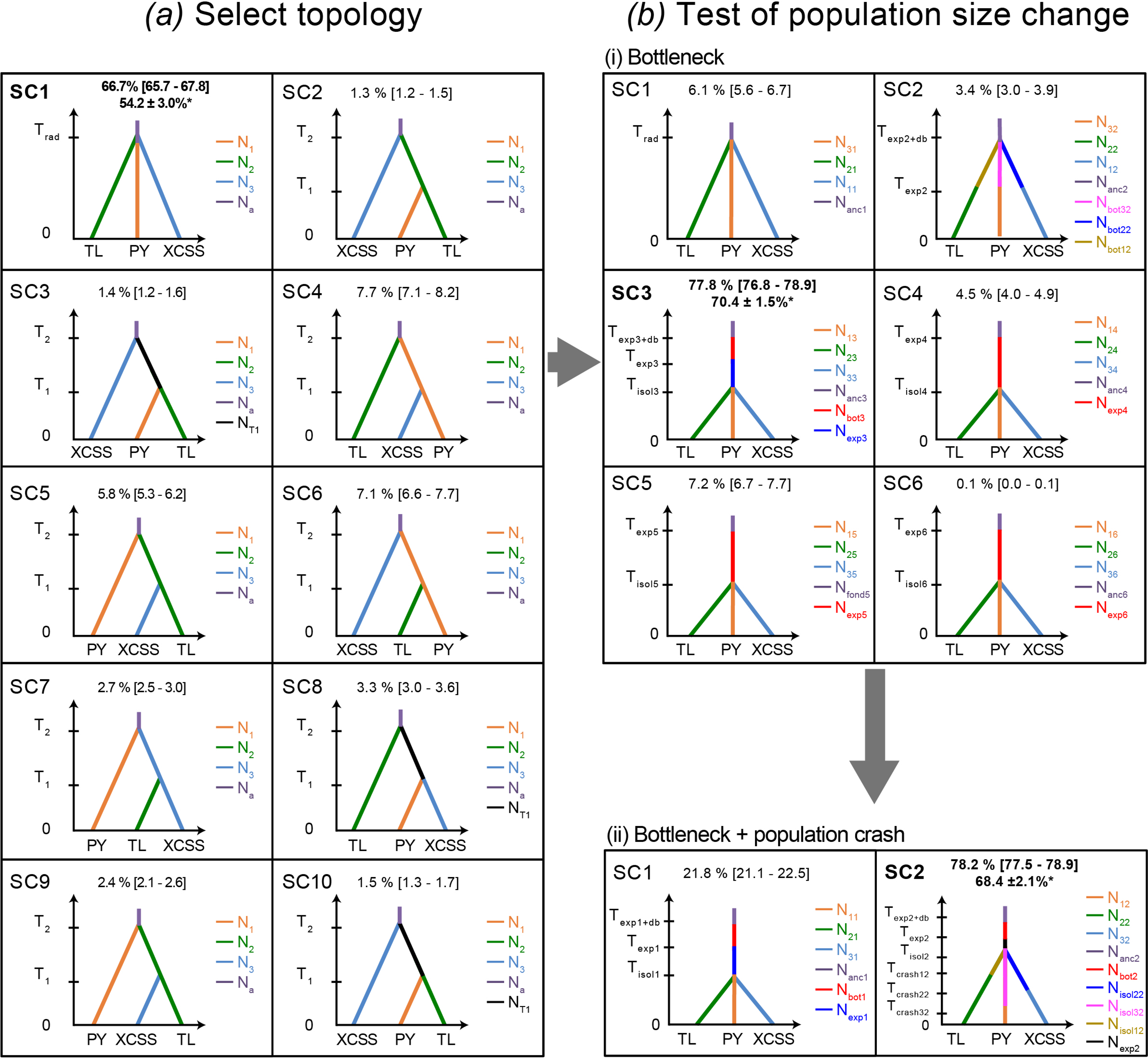
Schematic diagram of the ABC analysis to compare evolutionary histories and divergence scenarios generated and tested using the program DIYABC. Each coloured segment depicts a distinct effective population size. The posterior probability estimated using the ABC-LDA procedure is provided for each scenario. * indicates the posterior probability estimated using the ABC-RF is also provided for the best scenario of each step. See the main738 text, appendix S1, Table S2-S3 for further details.

The second ABC step tested for occurrence of simple changes in effective population size (*Ne*) during the divergence of the three populations under six nested competing scenarios (Fig. 4bi, Table S2 and S3). The scenario assuming a bottleneck in the ancestral population prior to population split (SC3) outcompeted the other five scenarios (*i.e.*, no change (SC1) or simple decline or expansion in the ancestral or daughter populations, SC2, SC4-6), with a probability of 78% and no overlap in 95%CI for the ABC-LDA, and 69.8 ± 1.9% under the ABC-RF (Fig. 4bi and Table S6). The sensitivity analysis showed that simulations generated with SC3 were more difficult to identify, with an average Type-I error rate of 12.4% (ranging between 1 and 21% error, depending on the scenario), and a total prior error rate of 62.7% under the ABC-LDA and 41.4% under the ABC-RF (Fig. S6 and Table S6). Nevertheless, the Type-II error rate (false positive) was only 7.2% and the power to discriminate this scenario from the others was 92.8%. Furthermore, the assessment of goodness-of-fit of this bottleneck scenario (SC3) to the data showed very good performance, as the simulations using this scenario and posterior distributions for each parameter, were able to reproduce all but one observed summary statistic, in contrast to all other scenarios (Table S6 and S7).

The final step (Fig. 4bii, Table S2 and S3) tested whether a recent demographic collapse in each population within the last 5 generations (or 50 years as reported in the literature^15^) could have produced a detectable genetic footprint. The scenario involving a recent collapse (SC2) in each population received a significantly higher probability (ABC-LDA 78.2%; ABC-RF: 69.2 ± 2.0%) compared with the alternative scenario of constant *Ne* since the population split (Fig. 4bii, Fig. S6, Table S8, and S9). Both Type-I and Type-II error rates were <16% indicating adequate power and sensitivity of our ABC analysis.

The model parameters estimated under the final best demographic scenario (SC2 in Fig. 4bii, Table S10 and S11) suggest that the ancestral population would have been large (*N_anc2_* = 18,700 individuals; 95%CI: [3,480 – 19,700]), and that a small fraction would have founded the Yangtze River populations ~3,400 95%CI: [1300 – 41,000] years ago (*T_exp2_*) and expanded to reach an effective size of about 5,660 individuals (*N_exp2_;* 95%CI: [2,900 – 9,840]). The three daughter populations would have then split from each other ~1,030 years ago (T_*isol2*_; 95%CI: [214 – 4,380]), and reached an effective size of about 2,000 individuals after their split. Each of these daughter populations would have gone through a significant decline leaving less than 2% of their pre-collapse size during the last 50 years (see Table 3, S10, and S11).

## Discussion

Our study shows that the present-day genetic diversity of finless porpoise in the Yangtze River (YFP) has been strongly influenced by an initial founder event that took place several thousand years ago, followed by a relatively recent split into 3 populations (XCSS, TL, PY), and a recent demographic collapse within the last 50 years. Indeed, consistent with previous studies^18,19^, the ABC genetic inferences showed that a few individuals coming from a large ancestral population, likely a marine population of *Neophocaena asiaeorientalis sunameri* in the Yellow Sea, colonized the Yangtze River within the last 41kyrs, and most likely during the Last Ice Age. The subsequent population split into three populations occurred between 200 and 5,000 years BP. This suggests that the population split was triggered before the intensification of human activities of the last 50 years in the Yangtze River. This event may thus have been related to the colonization process itself during the post-glacial period^19^ and/or other environmental or human-related factors. Consistent with the hypothesis of population split driven by post-glacial changes, episodes of contraction and expansion of the Yangtze River mainstream and the adjacent lakes occurred during the Holocene period^39^. Interestingly, the Yangtze River mainstream and the appendage lakes, including the Poyang and Dongting lakes, retracted and shrank significantly during the late Holocene (from *ca.* 3000 years BP to now)^39–41^. These environmental changes might have promoted the split of the ancestral population after the colonization of the Yangtze River.

Each population harboured a genetic footprint of dramatic population reduction. This especially affected the genetic diversity at the microsatellite markers which displayed very small values of the *M_GW_* statistic (Table 1) characteristic of significant recent decline^34^. This is also indicated by the ABC analysis supporting a scenario describing a drastic population reduction in each population within the last five generations (Fig. 4b_ii_, SC2). According to this scenario, the reduction in effective size would have been massive since their current sizes would be less than ~2% of their pre-collapsed sizes (Fig. 4, Table 3, S10 and S11). These genetic inferences are in line with the field estimates^14,15,42^, reporting a continuous decline of the YFP since the 1980s^10–15,43,44^. Half of the census populations in the main stem of the Yangtze River would have been lost within the past 15 years, with the abundance dropping from 2,500 porpoises in 1991^10^ to 1,225 in 2006^13^. With the 400 porpoises in the Poyang Lake and the 100 to 150 porpoises of the Dongting Lake, the total census size in the Yangtze River and the two adjacent lakes observed in 2006 was only 1,800 individuals. The most recent estimates from the YFDE2012 showed that the porpoises in the main stem of the Yangtze would have been reduced again by half with 505 individuals reported the middle and lower reaches of the Yangtze River and approximately 450 porpoises in PY, and 90 porpoises in DT^14^.

These very small census populations sizes (*N*) are consistent with the very small contemporary effective population size (*Ne*) estimated from genetic data (≤92 individuals) in each population, with the XCSS population being the smallest of all three differentiated populations (between 14 and 22 individuals, Table 3). Comparably low estimates have been reported in other cetacean species, such as the southern Iberian ecotype of harbour porpoise (*Phocoena phocoena meridionalis*) in European waters (Ne ≤ 80^17,45^) or the coastal ecotype of bottlenose dolphins (*Tursiops truncatus*) in European waters (Ne ≤ 77^46^). However only highly endangered populations, such as the Borneo Orangutans (*Pongo pygmaeus*)^47^ or the Canadian woodland caribou (*Rangifer tarandus*)^48^, have *Ne* values as low as those observed in the XCSS population. Extremely low *Ne* clearly translate the very low genetic diversity observed in each population of the YFP and imply very low numbers of breeding individuals in each populations of the Yangtze River and the adjacent lakes^49^. However, drawing a more direct link between *Ne* and *N* is actually difficult considering our study design. Previous studies have shown that no direct relationship can be expected between *Ne* and *N* when generations are overlapping, sampling spans several years, includes multiple cohorts and age classes, and when immigration may occur^50–52^.

The three YFP populations – XCSS, TL and PY– have not exchanged migrants over the last few generations according to our genetic estimates (Fig. 3 and Table S1). However, all three are or have been recently connected to the admixed group. We observed genetic evidence of unidirectional gene flow from the admixed group to XCSS and TL. This recent migration in the middle section of the Yangtze and XCSS in the upper section may be rare and/or no longer occurring based on the observed fixation of the mitochondrial haplotype in the XCSS population (Table 1 and Fig. 2d). This is further supported by the YFDE2006 and YFDE2012 surveys that reported increasing gaps in the distribution of the species in the upper section of the Yangtze River^14^. In contrast to the two other populations in the main stream river, gene flow was in the opposite direction in the Poyang Lake, from PY to the Admixed group (Fig. 3 and Table S1). This is consistent with field observations reporting groups of porpoises from PY moving to the main river stem in the morning and back to the lake in the afternoon^43^. This result also supports previous assertion^14^ that immigration of porpoises from PY to the river may dampen population decline in the Yangtze River. Unfortunately, such migration is likely insufficient given the observed ongoing decline^14,42^. In principle, the admixed group in the middle section of the Yangtze River could serve as a bridge connecting the three differentiated populations. However, our gene flow estimates do not support this (Fig. 3) and are consistent with the increasing observation of gaps in the distribution of the YFP and the loss of connectivity between populations.

Additional populations of YFP may exist in the Yangtze mainstream as the YFP is known to occur upstream and downstream from our study area (Fig. 1). For example, the three porpoises sampled around Shanghai city (SH) seem to belong to another differentiated unit (Fig. 2), but no definitive conclusions can be drawn at this time due to low sample size (n=3). Nevertheless, the present study provides a representative view of the population genetic structure, connectivity and demographic trends that can help to define priority areas where conservation measures need to be taken.

Drastic reduction in population abundance has left a clearly detectable genetic footprint on the genetic diversity of the YFP and coincides with the loss of connectivity between populations as well as the intensification of human activities along the Yangtze River over the last 50 years. To ensure that connectivity between populations is maintained, mitigation of human impacts need to include the entire river catchment^44^. For example, restricting fishing and sand-mining activities in the mouth area of Poyang Lake (Hukou, Fig. 1) could restore the lake-river migration of the YFP. Modification of current *in situ* reserves could improve connectivity between Ezhou and Zhenjiang (Fig. 1). Likewise, more active measures including a whole year fishing ban in the *in situ* natural reserves could certainly help^14^ and possible translocation of isolated individuals in the hope of increasing breeding opportunities could increase genetic diversity.

## Material & Methods

***Samples collection and DNA extraction*.** A total of 153 Yangtze finless porpoises were sampled between 1998 and 2011 across the distribution range (Fig. 1), including 3 from Shanghai (SH), 17 from Tongling (TL), 5 from Anqing (AQ), 15 from Ezhou-Huangshi (EZHS), 3 from Wuhan (WH), 14 from Honghu-Paizhou (HHPZ), 2 from Jianli (JL), 16 from Xingchang-Shishou (XCSS), 20 from Dongting Lake (DT) and 58 from Poyang Lake (PY). A detailed description of the sampling procedure and genomic DNA extraction is provided in Chen *et al.*^20^ Briefly, blood samples (n=113) were drawn from the caudal vein of live porpoises, immediately preserved in an Acid-Citrate-Dextrose solution, and stored in liquid nitrogen. Tissue samples (n=40, mainly muscle) were obtained from dead porpoises and preserved in 80% alcohol. Samples from the Yangtze mainstream and Dongting Lake were collected from accidentally killed or stranded individuals over the past decade. Fifty-eight blood samples from PY were collected from live animals during three field surveys conducted in early spring of 2009 (n=28), 2010 (n=17) and 2011 (n=13), under a special permit from the Poyang Lake Fishery and Fishing Administration Office of Jiangxi Province. Gender was identified by visual inspection of the genital parts. In total, 76 males and 77 females were sampled. The sampling was conducted in accordance with the Regulations of the People’s Republic of China for the Implementation of Wild Aquatic Animal Protection promulgated in 1993 by the Food and Agriculture Organization of the United Nations (FAO, FAOLEX No. LEX-FAOC011943, http://www.fao.org/faolex/results/details/en/?details=LEX-FAOC011943), adhering to all ethical guidelines and legal requirements in China.

Total genomic DNA was isolated from blood samples using the Whole Genome DNA Extraction Kit (SBS, Shanghai Inc.) following the manufacturer’s instructions. For tissue samples, total genomic DNA was extracted using a standard proteinase K digestion and phenol/chloroform extraction protocol^53^.

***Microsatellite and mitochondrial data set*.** Microsatellite and mitochondrial (mtDNA) data have been previously obtained and described by Chen *et al.*^20^ All 153 porpoises were genotyped at 11 polymorphic microsatellite loci (YFSSR1, YFSSR42, YFSSR59, YFSSR5, YFSSR40 from *N. p. asiaeorientalis*^54,55^, NP391, NP404, NP409, NP464, NP428 from *N. phocaenoides*^56,57^, and PPHO130 from *Phocoena phocoena*^58^. The genotyping protocol and quality checks have been described in Chen *et al.*^20^ In the subsequent analyses of the microsatellites, we only kept individuals for which we had at least 50% of the locus available (i.e. at least 5 microsatellite loci and the mtDNA or 6 microsatellite loci). Those filters excluded of 5 individuals, leading to a final data set of 148 individuals. The mtDNA data includes a 597 base-pairs fragment of the hyper-variable region 1 of the control region successfully sequenced for 129 individuals (see Chen *et al.*^20^ for details on the PCR and sequencing procedures). The microsatellite and mtDNA-CR data are available in Supplementary Dataset S1.

***Population genetic structure*.** We used the Bayesian model-based clustering of *STRUCTURE* v2.3.4^26^ to estimate the admixture proportions for each individual to each cluster identified in the microsatellite data. *STRUCTURE* cluster individual multilocus genotypes into K groups while minimizing departures from Hardy-Weinberg and Linkage Equilibria. We used the admixture *Locprior* model with correlated allele frequencies designed to detect weak signals of genetic structure without introducing bias or forcing the clustering^26^. We conducted a series of independent runs with different value for *K* from 1 to 5. Each run used 1×10^6^ iterations after a burn-in of length 2×10^5^. To assess that convergence of the Monte Carlo Markov Chains (MCMCs) had been reached, we performed 10 independent replicates for each *K* and checked the consistency of results using *CLUMPAK^59^.* We assessed which *K* value best fit with the data using (1) the likelihood of each K, following *STRUCTURE’s* user manual; (2) its rate of change with increasing K^60^; and (3) visual inspection of newly created clusters with increasing K^61^. Post-processing of the results, including generation of barplots, was conducted using *CLUMPAK*^59^. The geographic distribution of each group (Fig. 2) was mapped using the R statistical package *MARMAP* v.0.9.5^62^ and ETOPO dataset^63^.

The genetic structure in the microsatellite data was further inspected using a Discriminant Analysis of Principal Components (DAPC)^28^ and a Principal Component Analysis (PCA)^27,29^. These exploratory methods does not rely on any model assumptions and provides a complementary validation of the structure depicted by *STRUCTURE*^64^. These analysis were conducted using *adegenet* 2.0.1 package^65^ for R^66^ on centred genetic data (i.e. set to a mean allele frequency of zero), with missing data replaced by the mean, following the authors’ recommendations. The PCA was used to display the individual multilocus genotypes into a reduced multidimensional space defined by the first two principal components (PCs), colour-coding each individual according to the clustering identified by *STRUCTURE*, in order to assess the congruence. The DAPC method identifies genetic clusters by optimizing the difference between predefined groups and minimizing the variation within those groups^28^. It first reduces the number of variables using a PCA and then maximises the differences between groups using a Discriminant Analysis. The DAPC was performed with prior information on groups using the clusters defined by *STRUCTURE*. The number of PCs to retain and the reliability of the DAPC were determined using the cross-validation approach present in the *adegenet* 2.0.1 package^65^ for R^66^. As a result of this cross-validation step, a total of 60 PCs and 4 discriminant functions were retained to describe the relationship between the clusters. This number of PCs captured 80% of the total variation and provided the highest percent of correctly predicted subsamples with the lowest error. Finally, the score of each individual for the first two discriminant functions (DFs) were plotted as scatter plot in R and the memberships probability of each individual to the clusters defined by the DAPC were plotted as barplot in R.

***Genetic diversity and differentiation*.** We compared microsatellite genetic diversity between populations using the allelic richness (*A_r_*) and private *A_r_* (*pA_r_*) computed with *ADZE*^67^, and the observed and expected heterozygosity (*H_o_* and *H_e_*) computed with *GenAlEx* v6.5^68^. Departures from Hardy-Weinberg were tested using 10^4^ permutations with *FSTAT* v2.9.3.2^70^ and quantified using *F_IS_* and *F_ST_*^69^ in *GenAlEx* v6.5^68^. We applied a Bonferonni correction to correct for multiple tests.

Mitochondrial genetic diversity was estimated for each of the genetically distinct groups identified in the present study. Variation among sequences was measured using the number of segregating sites (*S*), number of singletons and shared polymorphisms, number of haplotypes, haplotype diversity (*H_d_*) and two estimators of population genetic diversity, *π* based on the average number of pairwise differences^71^ and *θ_W_* based on the number of polymorphic sites^72^. All statistics were calculated using *DNASP* v5.10.01^73^. We used the *F_ST_* statistics estimated from the average number of differences within and between populations^74^. Significance was tested with 1,000 permutations of Hudson’s nearest neighbour distance *Snn* statistics, which measures how often the nearest neighbour of a sequence (in sequence space) is from the same population^75^.

***Contemporary effective population sizes*.** We used *NeEstimator* v2.01^30^ as a first approach to estimate contemporary *Ne* based on linkage disequilibrium (LD) between loci, filtering out rare alleles with a frequency *P_crit_* ≤ 0.02 that could bias *Ne* estimate^76^. The second method implemented in *ONESAMP* v1.2^31^ also use LD as summary statistics among others in an Approximate Bayesian Computation (ABC) to estimate *Ne*, considering uniform prior between 2 and 500 of *Ne*.

***Contemporary gene flow between populations*.** We estimated contemporary effective migration rate (*m*) between populations using *BayesAss* v.3.0.3^32^. Preliminary runs were performed to adjust the mixing parameters of the MCMC and ensure proposal acceptance rates between 20% and 60% following authors’ recommendations. We then performed 12 independent runs with different seeds, a burn-in of 5×10^6^ iterations followed by 2×10^8^ iterations, and a sampling parameter values every 2000 iterations. Convergence of the MCMCs was checked by comparing the traces of each run using *Tracer* v1.5^77^ and by evaluating the Effective Sample Sizes (ESSs) of each parameter, keeping only runs where ESS ≥ 200. Model fitting to the data was assessed using the Bayesian Deviance Index using the R-script of Meirmans^33^. Runs that converged were combined to estimate the mean, median and 95% Highest Probability Density interval for each parameter in *Tracer* v1.5. The effective number migrants (*N_e_* × *m*) per generation between populations was obtained by combining *Ne* and *m* estimates.

***Genetic inference of population demographic history*.** We used *DIYABC* v2.1^78^ to estimate the *M_GW_* value and conduct 1×10^6^ coalescent simulations to produce a null distribution against which the observed value could be compared. The *P*-value indicates the proportion of simulations which provide a value below the observed one. For the mtDNA-CR, we used the Tajima’s *D*^35^ and tested for significant departure from a null expectations using 10,000 coalescent simulations in DNAsp^73^.

Next, we investigated the demographic history best describing the genetic diversity of the combined microsatellite and mtDNA markers using a coalescent-based ABC approach^22^. We subdivided our workflow in two nested parts (Fig. 4): identify the most likely population tree topology for our dataset among 10 plausible scenarios describing different potential population branching (Fig. 4a); and then test for evidence of changes in effective population size in the ancestral and daughter populations (Fig. 4b). This later part was further subdivided in two steps, (i) first testing for simple changes in effective population size (*Ne*) with six nested competing scenarios, and (ii) then testing whether adding the known population decline observed in each population over the last 50 yrs in the best scenario improved significantly the model fit to the data (Fig. 4b, Table S2 and S3).

For each part (Fig. 4), an ABC analysis was conducted using the program DIYABC v.2.1.0^78^ applying the following steps (Fig. S4): (1) coalescent simulations of 1×10^6^ pseudo-observed datasets (PODs) for each competing scenario and calculation of summary statistics (SS) describing the observed genetic variation for each POD; (2) select the best model by estimating the posterior probability (PPr) of each scenario using two approaches: the “standard” procedure relying on a logistic regression on 1% PODs producing SS values closest to the observed ones after a Linear Discriminant Analysis (ABC-LDA) as a pre-processing step^36^ and the recently introduced Random Forest (ABC-RF) procedure ^37,38^; (3) evaluate the confidence in scenario choice by estimating the type-I and type-II error rates based on simulated PODs using the ABC-LDA, as well as the prior error rate from the of the ABC-LDA and ABC-RF ^37,38^; (4) estimate the marginal posterior distribution of each parameter based on the best model including (among other) *Ne* and times of population size changes and splits (*T*); and finally, (5) evaluate the goodness-of-fit of the fitted model to the data. Details are provided in Fig. 4 and in the supplementary materials (Appendix S1, Fig. S4, Table S2, S3).

## Acknowledgments

We thank Pr. JL Olsen for constructive comments. This work was supported by the National Key Programme of Research and Development (2016YFC0503200) from Ministry of Science and Technology of China, the National Natural Science Foundation of China (Nos. 30730018, 31430080), the Special Fund for Agro-scientific Research in the Public Interest (No. 201203086), and the Special Conservation Fund for the Yangtze finless porpoise from the Ministry of Agriculture of China. This work was also supported by Major Project of Natural Science Research in Anhui Province (No. KJ2016A863).

## Author contributions

MCF, JZ, DW designed the study; MC, JZ, ZM, YH, KW, MW, QZ, DW conducted the field expeditions and collected the samples; MC conducted the laboratory experiments and collected the data; MCF, YBC and FL analysed the data; MCF interpreted the results and wrote the manuscript with help from YBC, MC, JZ and DW, and final approval by all co-authors.

## Additional information

***Data accessibility*.** All data generated during this study are included in this article (and its Supplementary Information files).

***Competing financial interests*:** the authors declare no competing financial interests.

**Supplementary information** accompanies this paper at http://www.nature.com/srep

